# Short-wavelength-sensitive 2 (Sws2) visual photopigment models combined with atomistic molecular simulations to predict spectral peaks of absorbance

**DOI:** 10.1101/2020.08.13.249755

**Authors:** Dharmeshkumar Patel, Jonathan E. Barnes, Wayne I. L. Davies, Deborah L. Stenkamp, Jagdish Suresh Patel

## Abstract

For many species, vision is one of the most important sensory modalities for mediating essential tasks that include navigation, predation and foraging, predator avoidance, and numerous social behaviors. The vertebrate visual process begins when photons of the light interact with rod and cone photoreceptors that are present in the neural retina. Vertebrate visual photopigments are housed within these photoreceptor cells and are sensitive to a wide range of wavelengths that peak within the light spectrum, the latter of which is a function of the type of chromophore used and how it interacts with specific amino acid residues found within the opsin protein sequence. Minor differences in the amino acid sequences of the opsins are known to lead to large differences in the spectral peak of absorbance (i.e. the λ_max_ value). In our prior studies, we developed a new approach that combined homology modeling and molecular dynamics simulations to gather structural information associated with chromophore conformation, then used it to generate statistical models for the accurate prediction of λ_max_ values for photopigments derived from Rh1 and Rh2 amino acid sequences. In the present study, we test our novel approach to predict the λ_max_ of phylogenetically distant Sws2 cone opsins. To build a model that can predict the λ_max_ using our approach presented in our prior studies, we selected a spectrally-diverse set of 11 teleost Sws2 photopigments for which both amino acid sequence information and experimentally measured λ_max_ values are known. The final first-order regression model, consisting of three terms associated with chromophore conformation, was sufficient to predict the λ_max_ of Sws2 photopigments with high accuracy. This study further highlights the breadth of our approach in reliably predicting λ_max_ values of Sws2 cone photopigments, evolutionary-more distant from template bovine RH1, and provided mechanistic insights into the role of known spectral tuning sites.

**Author Summary:** In vertebrates, color vision depends on the complement of cone visual photopigments that have different spectral peaks of absorbance (λ_max_) within the cone population. Together, the type of chromophore and the amino acid sequence of the opsin protein directly affect the λ_max_ value. To understand this relationship further at a structural level, we previously developed a new molecular modeling approach to study Rh1 and Rh2 opsin classes by combining homology modeling, molecular dynamics simulations to extract structural parameters of chromophore conformations and statistical modeling. Here, we used this novel modeling approach to accurately predict the λ_max_ values for teleost Sws2 photopigments. Such a genome-to-phenome approach for predicting visual pigment function will be of great interest to evolutionary biologists, vision scientists, and molecular modelers, to better understand the diversity and mechanisms of sensory function. Moreover, it will pave the way for novel strategies to forward engineer visual pigments suitable for optogenetics applications.

## Introduction

For many animals, vision is a critical sensory modality that facilitates essential tasks that include navigation, predation and foraging, predator avoidance, and numerous social behaviors. In vertebrates, vision is initiated when photons enter the eye and interact with rod and cone photoreceptors found within the neural retina. Specifically, light is detected by photosensitive visual photopigments that are housed within the folded membrane of photoreceptor outer segments. Upon the absorbance of photons, these photopigments activate a specific phototransduction cascade that results in a change in membrane potential. Once hyperpolarized, visual photoreceptors cause a change in neurotransmitter release that propagates neural signals through other retinal neurons that ultimately lead the visual centers of the brain [1–3].

A vertebrate visual photopigment consists of a transmembrane opsin protein that is covalently linked to a vitamin A-derived chromophore. Indeed, it is the interaction of the chromophore with specific amino acids of the protein sequence of the opsin that results in a broad array of different spectral peaks of absorbance (i.e. the λ_max_ value) [1–6]. In the visual system, the predominant chromophores are either based on 11-*cis* retinal (i.e. rhodopsins or vitamin-A_1_ photopigments) or 11-*cis*-3,4-didehydro retinal (i.e. porphyropsins or vitamin-A_2_ photopigments) [1–3,7]. Throughout vertebrate evolution, the associated visual opsin genes and their gene products have diverged into five classes: these comprise a long-wavelength-sensitive (*LWS*) opsin gene class, two short-wavelength-sensitive (*SWS*) opsin gene classes (*SWS1* and *SWS2*), and two medium-wavelength-sensitive (*mws*) opsin gene classes called rhodopsin-like 1 (*RH1*) and rhodopsin-like 2 (*RH2*), respectively [1–3,8]. In general, RH1 opsins form the highly sensitive visual photopigments (rod opsins) of rod photoreceptors, which typically mediate dim light or scotopic vision by maximally detecting wavelengths at around 500 nm [1–3], although λ_max_ values can shift to shorter wavelengths (e.g. 470-490 nm in some deep-sea fishes [9–11]. The remaining four opsin gene classes (*LWS*, *SWS1*, *SWS2* and *RH2*) form the visual photopigments expressed in cone photoreceptors that mediate bright light or photopic vision. Color vision is possible when at least two cone photopigments with distinct λ_max_ values and overlapping spectral absorbance profiles are present within distinct cone populations, thus providing differential input to other retinal neurons [12]. Cone opsins of the SWS1 class typically produce photopigments with λ_max_ values between 360 nm and 450 nm (i.e. perceived as ultraviolet (UV) to violet parts of the light spectrum), SWS2 photopigments present with λ_max_ values of ~400-470 nm (perceived as blue), RH2 opsins produce photopigments with λ_max_ values of ~480-530 nm (perceived as blue-green), whereas LWS photopigments have λ_max_ values that are largely sensitive to a range of wavelengths from 500-570 nm (perceived as red) [1–3,8]. These spectral ranges are based on photopigments that possess a vitamin-A_1_-derived chromophore (i.e. rhodopsins); whereas for porphyropsins, the presence of a retinal chromophore based on vitamin-A_2_ shifts the λ_max_ value towards longer wavelengths (e.g. up to ~620 nm [1–3,7,13], a spectral property that is more pronounced the longer the wavelength [14].

There is a great deal of diversity within each class of vertebrate opsins, since visual photopigment proteins are under strong natural selection [15–17]. This diversity is particularly striking within the teleost fishes, where many of the opsin genes have been tandemly or otherwise replicated, followed by subfunctionalization or neofunctionalization to generate new photopigments with distinct λ_max_ values [10,18–21]. For example, many fish genomes harbor two or three copies of the *sws2* cone opsin gene [22], the subject of the present study. In some cases, only minor changes in the amino acid sequence have resulted in major changes in the λ_max_ value, as was demonstrated for the A269T (Amino acid numbering in this study is based upon bovine RH1 sequence numbering; S1 Fig) change in the Sws2a opsin of the spotted flounder (*Verasper variegatus*) that resulted in a photopigment that is sensitive to longer wavelengths (i.e. λ_max_ = 485 nm instead of 466 nm) [23].

There is great interest in understanding the evolutionary, as well as the molecular mechanisms, that underlie the diversity of visual photopigments and their spectral peak absorbances. In general, there are two main experimental techniques for defining the λ_max_ value of visual photopigments, namely microspectrophotometry, which analyzes the spectral profile of photoreceptors directly, but only with fresh retinas, and spectral tuning site substitutions cannot be studied in isolation [24] or *in vitro* regeneration [2,23,25–27]. The latter is a popular, yet highly specialized and labor-intensive, approach that has been used to deduce the spectral properties of photopigments in isolation from wildtype and inferred ancestral protein sequences, followed by reconstitution with chromophore (usually 11-*cis* retinal) and experimental measurement of absorbance from 200-800 nm [25,28–31]. This technique, when combined with site-directed mutagenesis, has illuminated the contributions of specific amino acid substitutions to shifts in the λ_max_ value [2,25,26,30]. Although success of such studies has permitted the accurate prediction of the spectral peak of absorbance for some opsin classes (specifically, LWS [32] and UV-sensitive SWS1 [26] photopigments), such efforts are far from sufficient to understand pigment function that allows prediction of the λ_max_ value based entirely upon the amino acid sequence [6,33]; this is particularly the case for both SWS2 and RH2 photopigments, where experimental interventions are frequently employed. One of the long-term goals of our studies is to develop computational tools that result in straightforward, genome-to-phenome, predictive pipelines, as is the case for the application of spectral modeling and atomistic molecular simulations for both the Rh1 and Rh2 classes of visual photopigments [10,34].

The λ_max_ value of any functional photopigment in its inactive form is determined by the conformation adopted by the chromophore in the dark state, a function that is dependent upon the shape and composition of the retinal binding pocket, as well as the counterions that stabilize the Schiff base linkage of the chromophore to lysine (K) 296 of the opsin protein [35–37]. Therefore, our aim of generating genome-to-phenome pipelines for predicting the λ_max_ values of visual photopigments from their amino acid sequences, has been to increase an understanding of chromophore conformation via atomistic molecular simulations and to use this structural information to generate predictive models [10,34]. The approach includes: 1) building homology models for classes of visual photopigments, using the solved crystal structure of bovine rod opsin (RH1) as a template [38]; 2) carrying out atomistic molecular dynamics (MD) simulations using homology models of photopigments with experimentally-measured λ_max_ values; 3) identifying structural features of the chromophore and opsin that are correlated with a particular λ_max_ value; and 4) using these features to generate a statistical model that can in turn be used to predict λ_max_ values of other photopigments. This approach was successful for predicting λ_max_ values of teleost Rh1 photopigments [10] and a closely-related Rh2 class of teleost cone photopigments [34]. Notably, this approach also revealed structural features of the Rh1 protein, namely the presence vs. the absence of a C111-C188 disulfide bridge that powerfully predicts λ_max_ values >475 nm when present vs. λ_max_ values <475 nm when absent [10].

In this present study, we test the hypothesis that the approach outlined above and in our previous publications [10,34], can also successfully predict λ_max_ values of a class of cone photopigments that are phylogenetically-distinct from the known bovine RH1 template, namely the teleost Sws2 class. The SWS2 opsins are more divergent from RH1 than RH2 vs. RH1 [8,21] and are known to be notoriously difficult when attempting to successfully predict λ_max_ values from the amino acid sequence alone [1,2]. Furthermore, teleost Sws2 opsins display only ~48-51% amino acid sequence identity to the bovine rod opsin sequence (Table 1). By contrast, our previous studies tested this approach for teleost Rh1 photopigments, with ~49-83% identity to bovine rod opsin [10], and Rh2 cone pigments, with ~63-72% identity to bovine rod opsin [34]. Here we also test the hypothesis that this approach will work for a class of photopigments with a broader, and more short-wavelength-shifted range of λ_max_ values (i.e. 397-485 nm; Table 1) than the Rh1 (444-519 nm) or Rh2 (467-528 nm) photopigments used in our prior studies [10,34].

**Table 1.**
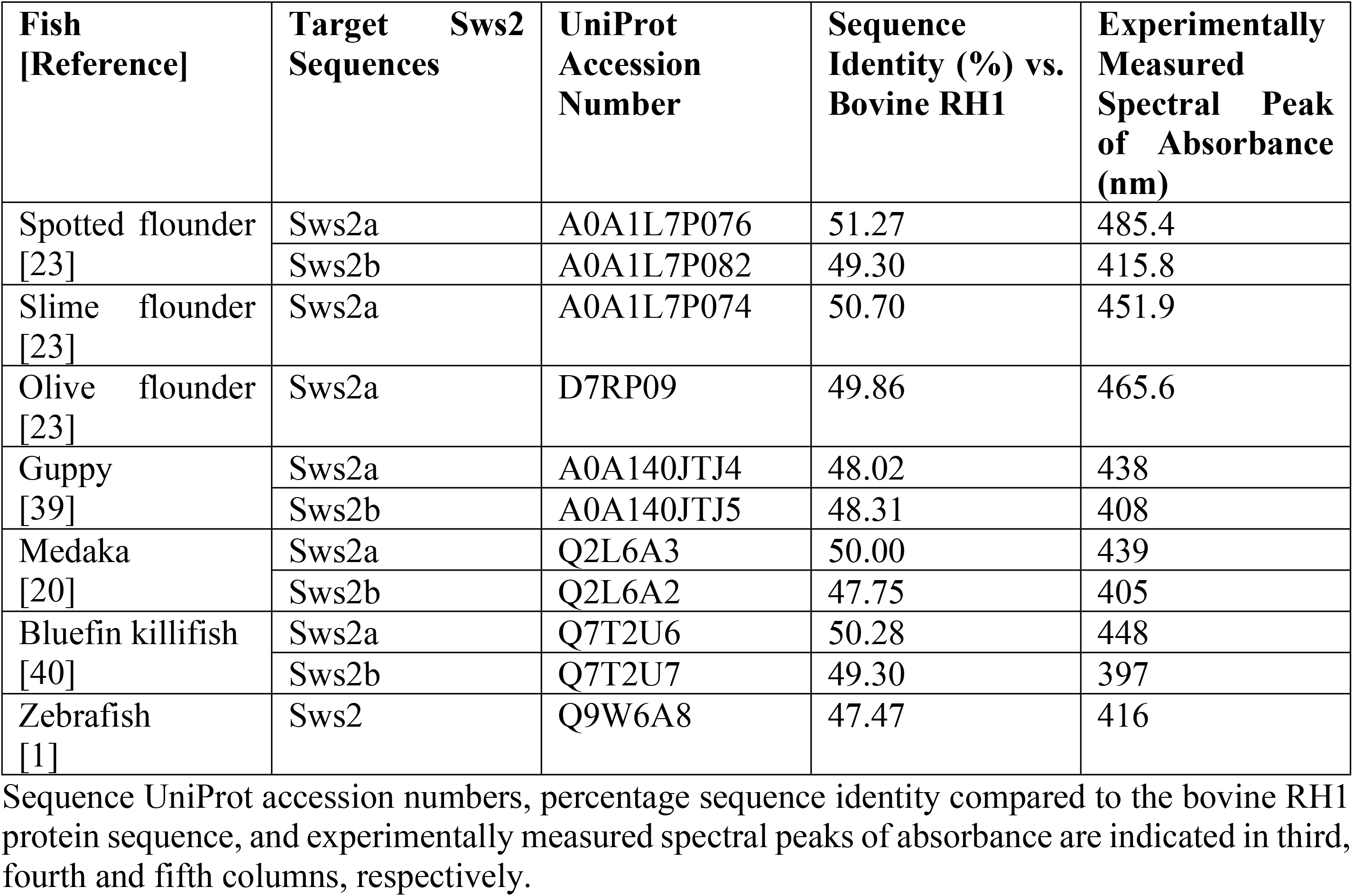
Selected Sws2 opsin sequences used for homology modeling.

In this study, we show that this approach was highly successful at predicting spectral peaks of absorbance values of 11 teleost Sws2 opsins for which sequences and λ_max_ values are known (Fig 1, Table 1). We identified three parameters of chromophore conformation that together accurately predict the λ_max_ value. Furthermore, we discuss these results in the context of known amino acid substitutions that likely contribute to divergent λ_max_ values [22]. These studies, therefore, not only provide a valuable extension of our prior work, but also guidance and strategic directions for the improvement of functionally-predictive photopigment modeling.

**Fig 1.**
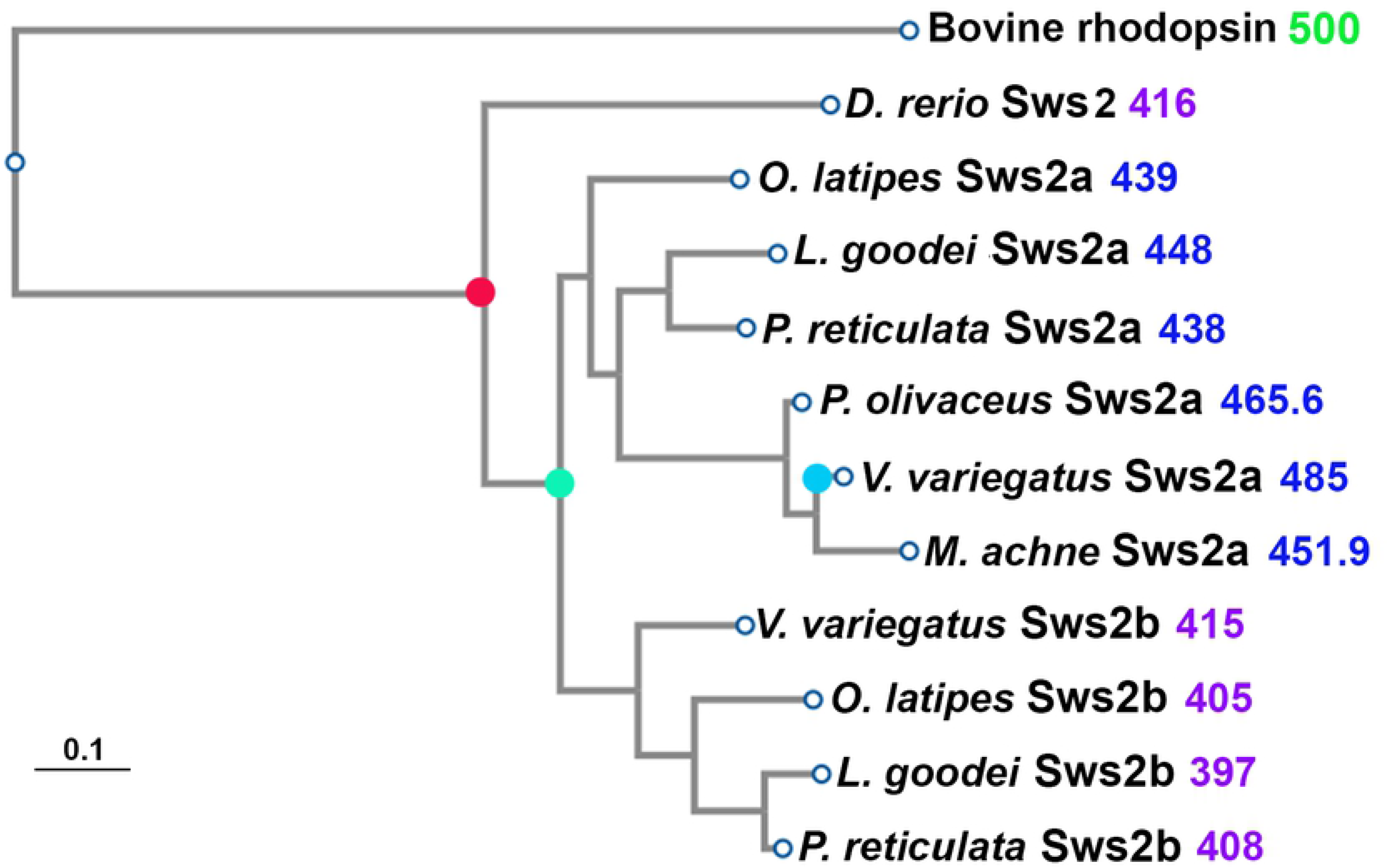
Evolutionary relationships of teleost Sws2 opsin proteins used in the simulations, as inferred by PhyML. Red filled circle indicates the speciation event that occurred prior to the duplication (green filled circle) of the *sws2* opsin genes in teleosts [18]. The duplication generated the *sws2a* clade, which encode photopigments with λ_max_ values that are shifted to longer wavelengths, and the *sws2b* clade, which encode photopigments with short-wavelength-shifted λ_max_ values. The blue filled circle indicates the amino acid substitution A269T (with numbering standardized to the bovine rod opsin sequence) that is likely to be the spectral tuning site important for a further shift of the λ_max_ value of *V. variegatus* Sws2a to longer wavelengths [23]. Experimentally measured λ_max_ values (in nm) are indicated next to the name of each opsin and are color-coded for λ_max_ values that are <430 nm (violet) or >430 nm (blue).

## Results

To develop a model to predict spectral peaks of absorbance (i.e. λ_max_ values) from a diverse set of teleost Sws2 cone opsins, the following amino acid sequences were selected: two from *V. variegatus* (spotted flounder) [23], one from *Microstomus achne* (slime flounder) [23], one from *Paralichthys olivaceus* (olive flounder) [23], two from *Poecilia reticulata* (guppy) [39], two from *Oryzias latipes* (medaka) [20], two from *Lucania goodei* (bluefin killifish) [40], and one from *Danio rerio* (zebrafish) [1]. These Sws2 opsin amino acid sequences show a wide range of experimentally measured λ_max_ values that range from 397 nm to 485 nm (Table 1). Such functional divergence probably evolved as primary *sws2* gene sequences mutated and were positively conserved, which likely led to distinct conformations of the Sws2-associated chromophore, 11-*cis* retinal, in the dark state (Fig 2).

**Fig 2.**
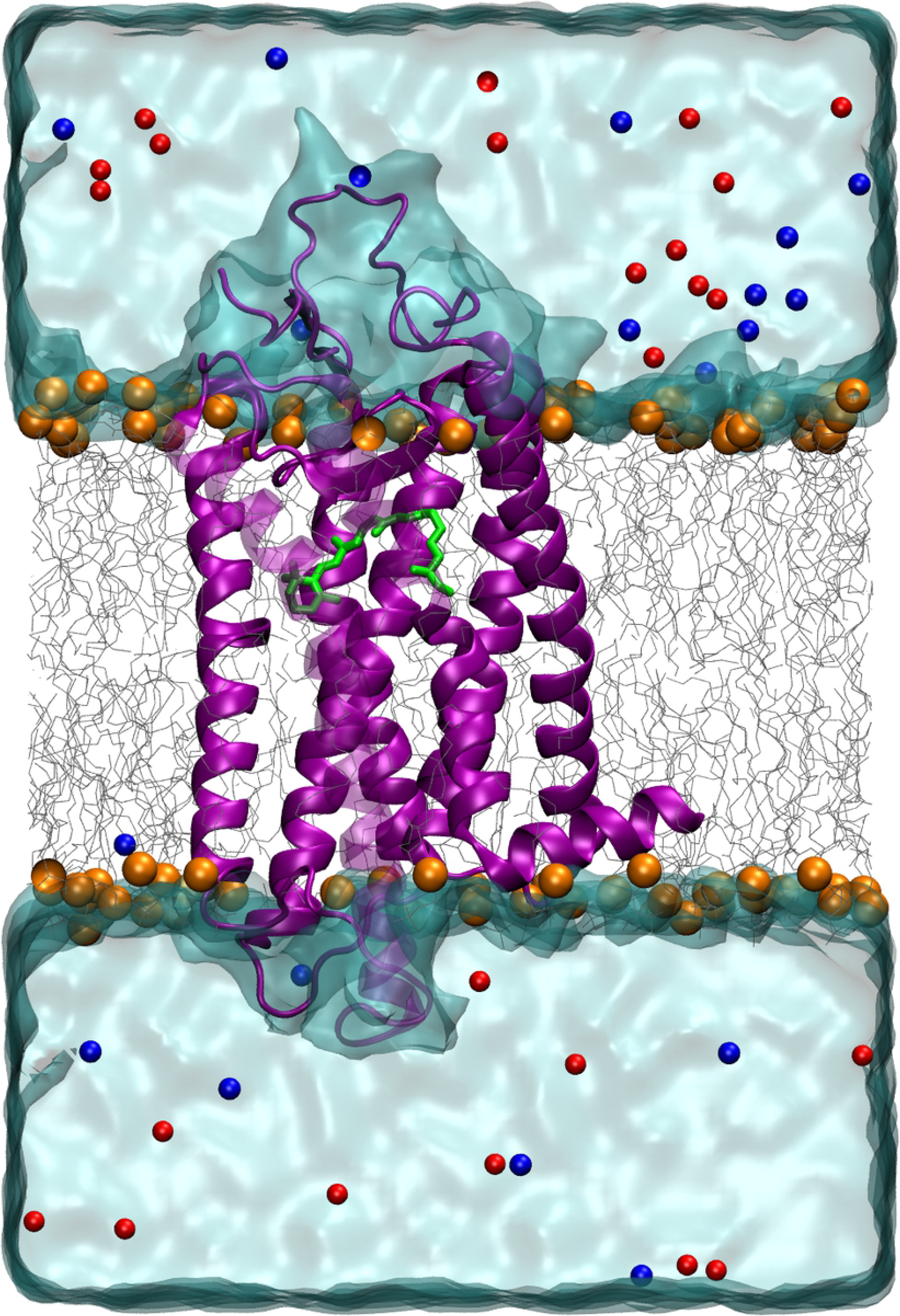
A representative 3D structure of Sws2 cone opsin (λ_max_ values <430 nm) homology structure (violet) with the chromophore (green) bound covalently to K296 of the opsin protein. It is inserted in a phospholipid bilayer (gray, carbon atoms; orange, phosphorus atoms) and surrounded by water molecules (light blue). Blue and red spheres indicate positive and negative counter ions, respectively.

We used a similar protocol that was shown to be successful for the prediction of Rh1 and Rh2 λ_max_ values [10,34], through building homology models using bovine rod opsin (RH1) as the template [41]. RH1 opsins are primarily present in vertebrate rods [8] and are the only mammalian visual pigment class where the protein structure has been experimentally determined [38,41]. Therefore, this, and in particular the bovine rod opsin protein structure, is the only template available for accurate homology modeling of vertebrate cone opsins. The sequence identity of the bovine RH1 template compared to teleost Rh1 rod opsins and Rh2 cone opsins ranged from 49-83% and 63-72%, respectively; which proved to be more than sufficient for the reliable modeling and accurate prediction of λ_max_ values [10,34]. In the present study, we test whether this approach is also reliable for predicting λ_max_ values of more evolutionarily-distant Sws2 cone opsins (Fig 1) that have ~48-51% sequence identity to the bovine RH1 template (Table 1). Furthermore, the selected Sws2 cone photopigments display a broader range and blue-wavelength-shifted λ_max_ values (i.e. 397-485 nm) compared to either Rh2 cone photopigments or rod opsins presented in prior teleost studies: 467-528 nm for Rh2 [34] and 444-519 nm for Rh1 [10] compared to the bovine RH1 template with a λ_max_ value at 498 nm [9]. Therefore, this present study also tests the capacity of our prior approach of combining homology modeling and MD simulations to accurately predict λ_max_ values for an opsin class that is difficult to predict from just the amino acid sequences, that are short-wavelength-shifted compared to the bovine template, and cover 88 nm of the electromagnetic spectrum.

Using the sequence information of 11 teleost Sws2 opsins, we built homology models based on the bovine rod photopigment (RH1 opsin + 11-*cis* retinal chromophore) structure as a template (Protein Data Bank (PDB) ID: 1U19) [41]. These homology structures with the chromophore attached covalently to K296 within the binding pocket were placed in lipid bilayers and water models (Fig 2). Each of these systems were then subjected to 100 ns classical MD [42] simulations using the protocol described in the Methods section (S1 Movie).

We analyzed the MD simulations for all 11 visual photopigments to understand the dynamics and to identify structural features associated with the chromophore and attached lysine residue (Fig 3A) that could potentially be used to explain differences in the λ_max_ values of the representative Sws2 photopigments used in this study. To understand the dynamics of the chromophore within the opsin binding pocket, we visualized the conformations of the chromophore seen in violet-(λ_max_ values <430 nm) vs. blue-sensitive (λ_max_ values >430 nm) photopigments (Fig 3B), where we observed a relatively compact cluster of chromophore conformations for blue-sensitive photopigments compared to violet-sensitive photopigments. This difference was more evident for the β-ionone ring and two methyl groups present at positions C9 and C13. In our previous study, the area under the curve (AUC) of root mean square fluctuations (RMSF) served as an additional feature of the chromophore that helped to predict the λ_max_ values of Rh2 cone visual photopigments [34]. We, therefore, first calculated RMSF values of all the heavy atoms of the chromophore and the linked lysine residue (LYS+RET) (Fig 3C) for each Sws2 photopigment as follows:

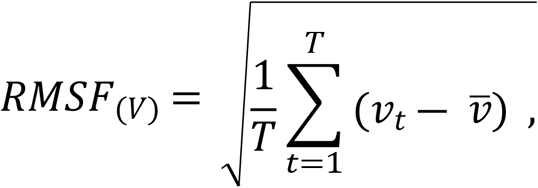

where T is the total number of molecular dynamics trajectory frames (V) and then calculated AUC of RMSF values (AUC RMSF_(LYS+RET)_) for each Sws2 photopigment. The atoms within the blue-sensitive photopigments clearly show lower RMSF values compared to the violet-sensitive photopigments (Fig 3C), suggesting that this chromophore feature may also be useful in predicting of λ_max_ values. Interestingly, this outcome is the opposite to what one would anticipate based upon our prior study, in which lower values of AUC RMSF_(LYS+RET)_ were associated with photopigments with shorter wavelength λ_max_ values [34].

**Fig 3.**
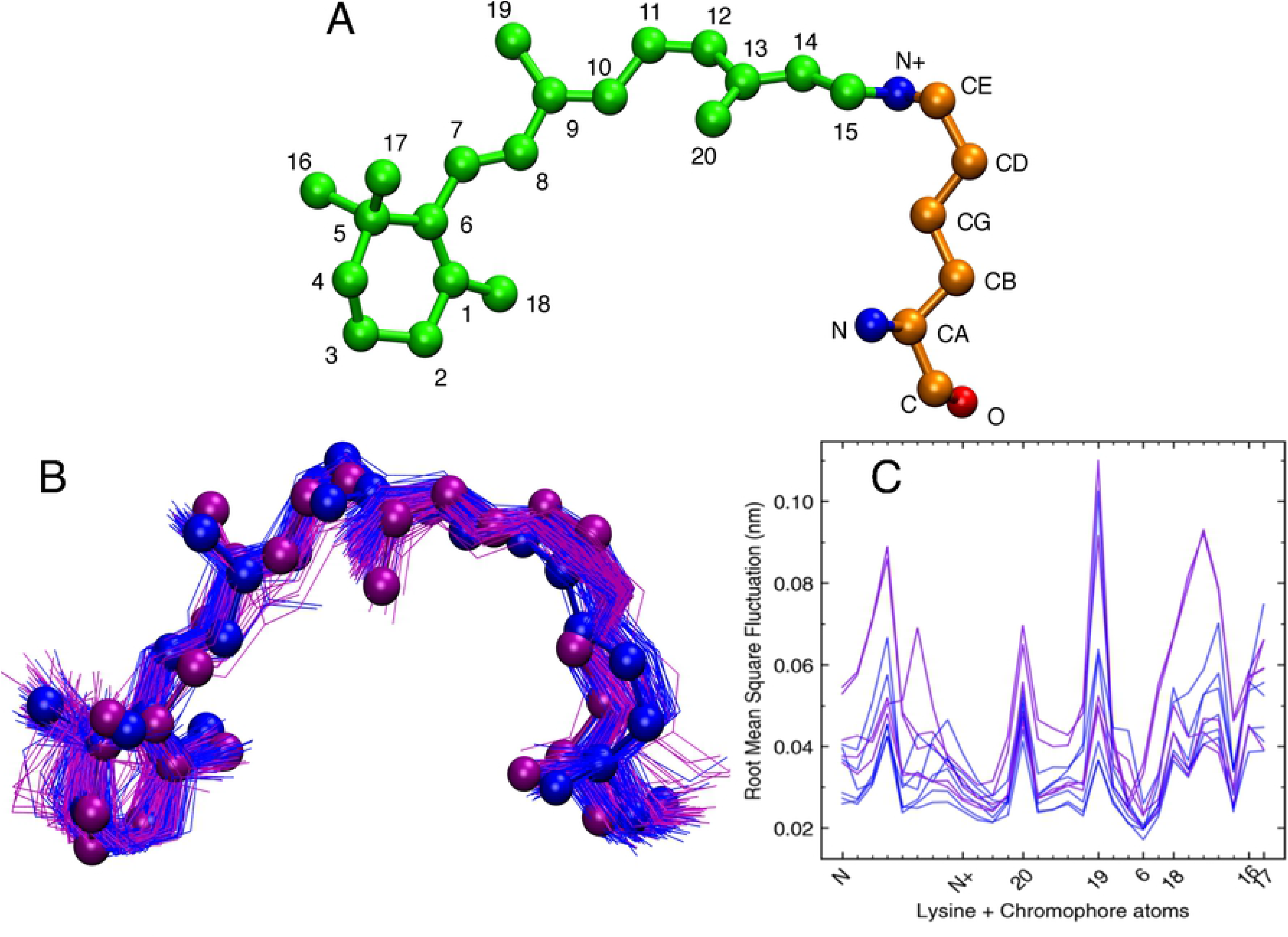
Conformations and fluctuations of 11- *cis* retinal chromophore and attached lysine in Sws2 photopigments. A) 3D orientation of 11-*cis* retinal linked to K296 of the opsin protein. B) Superposition of 11-*cis* retinal conformations from MD simulations trajectories. C) Root mean square fluctuation (RMSF) of 11-*cis* retinal linked to K296 (LYS+RET). The horizontal axis represents atoms of the LYS+RET. Blue-vs. violet-colored ball-and-stick conformations are those associated with Sws2 photopigments with λ_max_ values >430 nm vs. λ_max_ values <430 nm, respectively.

The dynamic nature of the chromophore during MD simulations suggests that we may use the geometric angles, dihedrals and AUC RMSF_(LYS+RET)_ parameters to differentiate between blue-sensitive and violet-sensitive Sws2 photopigments. For each photopigment, we examined a total of 19 angles (15 Torsion Angles and 4 Geometric Angles) (S2 Fig, Fig 3A) formed by the heavy atoms of the lysine residue at position 296 of the opsin covalently linked to 11-*cis* retinal. The model parameters that showed a reasonable correlation to experimental λ_max_ values were the median values of Torsions 3, 9, 10, 11, 12, as well as Geometric Angles 1 and 3, from a total of 19 examined angles. The standard model selection procedure was then used to determine the simplest linear regression model that best fitted to the parameters showing a reasonable correlation to experimental λ_max_ values. From our model selection procedure, we found the simplest model for the 11 Sws2 photopigments examined contained three terms: the median values of Torsion 3 (C15–C14–C13–C20), Torsion 12 (C19–C9–C8–C7), and Angle 3 (C3–C7–C8) (Fig 4). AUC RMSF_(LYS+RET)_ values were not identified by the model selection procedure as being predictively useful.

**Fig 4.**
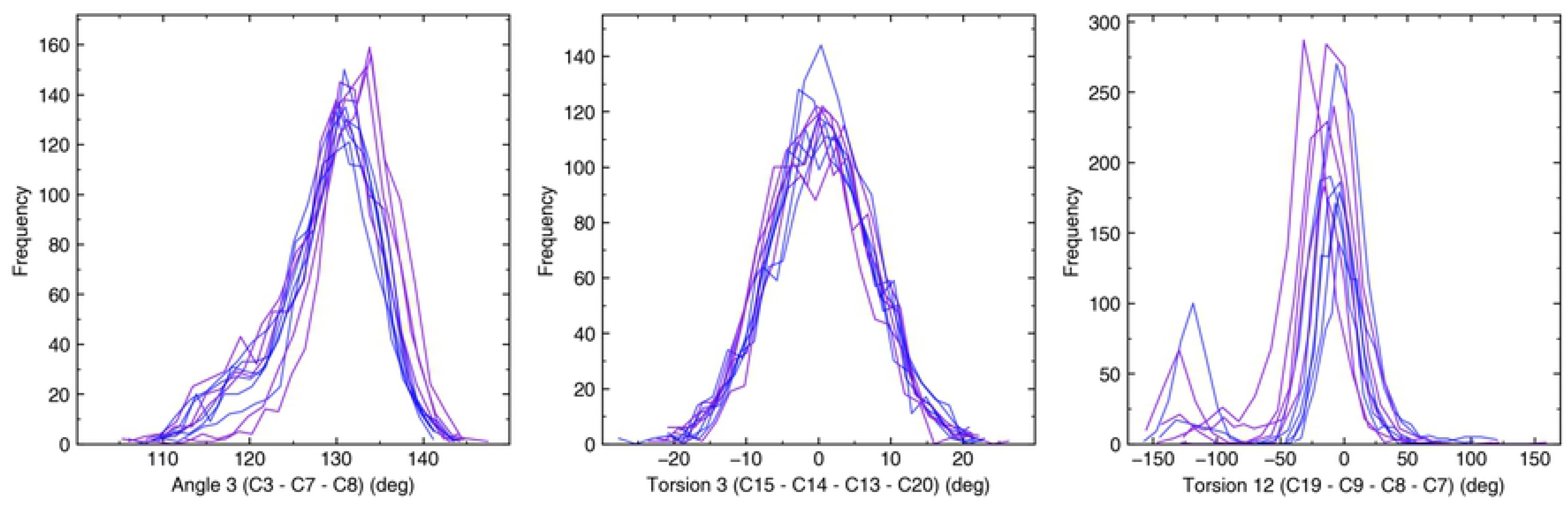
Frequency distribution of Angle 3, Torsion 3 and Torsion 12 observed in each opsin simulation. Blue and violet colors correspond to Sws2 photopigments with λ_max_ values >430 nm vs. λ_max_ values <430 nm, respectively.

The full model is explicitly given by:

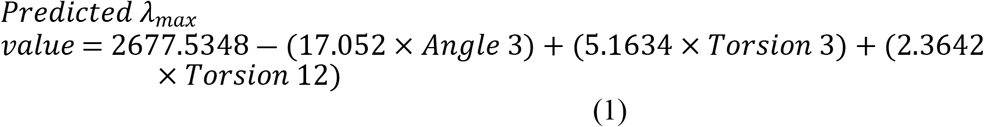

The larger values of Angle 3 predicted a spectral shift towards shorter wavelengths (i.e. a violet shift), while larger values of Torsion 3 and Torsion 12 predicted a shift towards longer wavelengths (i.e. a blue shift). Fig 5 shows empirically determined λ_max_ values vs. the model predicted values for each Sws2 photopigment analyzed, where our full model highly correlates with experimental data (R^2^ = 0.95). The error in prediction (error = | pred − exp |) ranged from 1.21 nm to 8.35 nm with an average error of 5.44 nm. To further test our statistical model, we carried out a “leave-one-out” analysis, where each Sws2 photopigment was removed from the regression analysis to obtain the coefficients for a model using Angle 3, Torsion 3 and Torsion 12 parameters, and then the λ_max_ value of the removed photopigment was predicted based upon the new linear model. The correlation of the individual predictions based upon only 10 pigments was reduced, but it was still highly acceptable (R^2^ = 0.86) (Fig 5). The lower correlation coefficient derived from the “leave-one-out” approach is largely explained by the less accurate prediction of the λ_max_ value for the zebrafish Sws2 photopigment (i.e. experimental λ_max_ value at 416 nm vs. “leave-one-out” predicted λ_max_ value at 400 nm). This less accurate prediction is likely due to Torsion 3, which has a relatively higher median value compared to Torsion 12 with a relatively lower median value for zebrafish compared to the median values of Torsion 3 and Torsion 12 of Sws2 pigments from other species. Nevertheless, our results strongly indicate that the statistical model derived from MD simulations of predicted Sws2 visual photopigment structures has the power to accurately predict their λ_max_ values.

**Fig 5.**
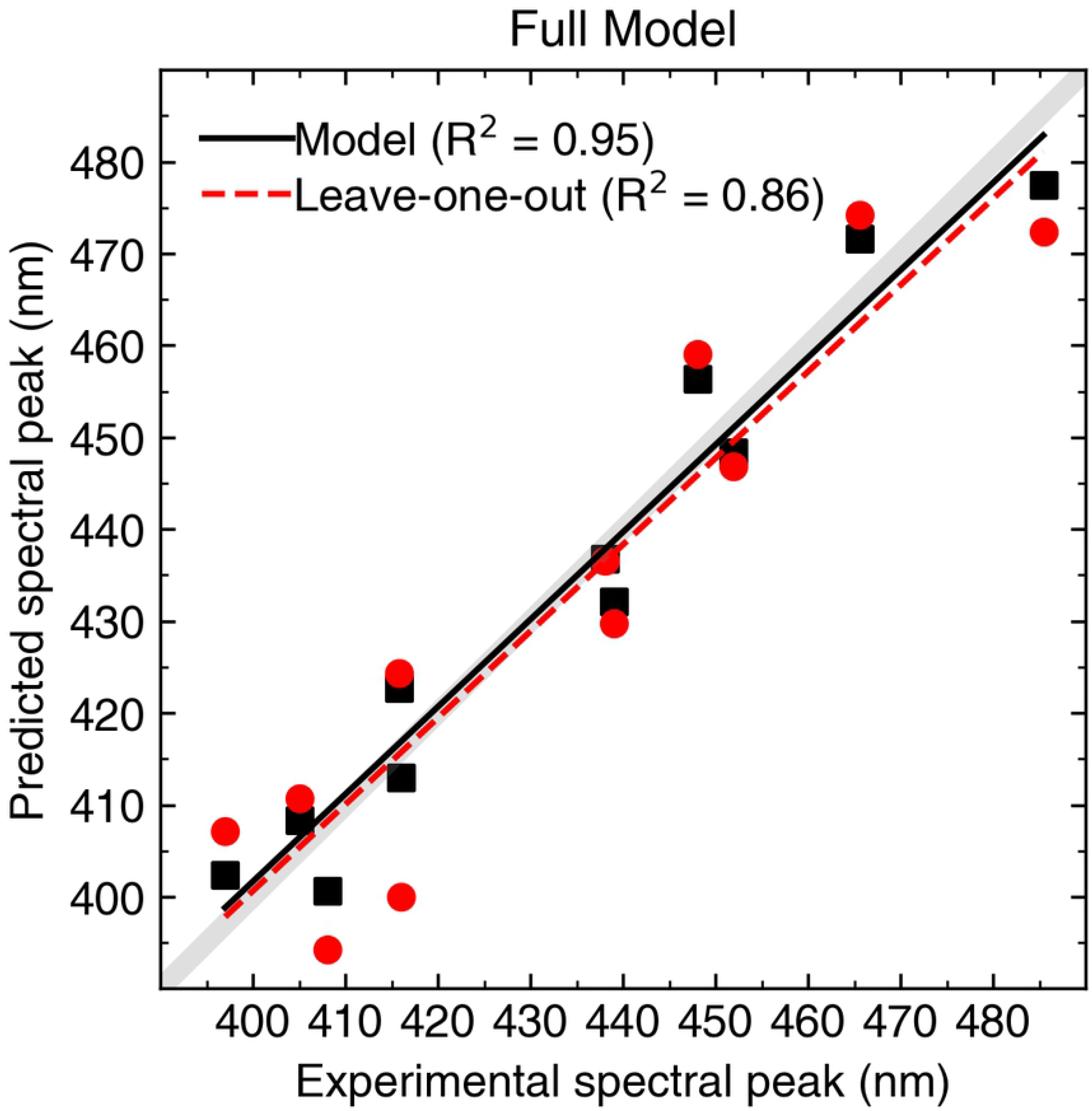
Experimental spectral peaks of absorbance (λ_max_) compared to predicted λ_max_ values by the full model equation 1 outlined in the main text for all 11 Sws2 photopigments analyzed. Gray lines indicate a perfect (100%) correlation. Solid black lines and black symbols represent the linear relationships between model-predicted and the experimental λ_max_ values, whereas dashed red lines and red symbols show linear relationships between “leave-one-out” predictions and experimental λ_max_ values. Corresponding correlation coefficients for both approaches are indicated.

To further test this approach and to simulate performance at predicting unknown λ_max_ values for Sws2 photopigments, we performed a “leave-one-out” approach at a species level. For each species where both Sws2a and Sws2b opsins are represented in the dataset (Table 1), a subset was generated where their Sws2 chromophore parameters were removed. A new model based on the newly generated subset was chosen using the full model selection procedure detailed in the Methods section. This new best-fit model was then used to predict the spectral peaks of absorbance for the omitted opsins. In this study, there are four species with both Sws2a and Sws2b photopigments (Table 1), namely the medaka, the bluefin killifish, the guppy, and the spotted flounder. For these four species, the following models were determined:
 
i. With the medaka Sws2 photopigment removed:

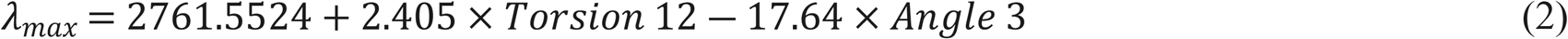
ii. With the spotted flounder Sws2 photopigment removed:

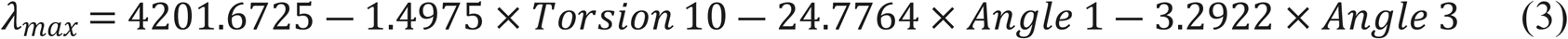
iii. With the bluefin killifish Sws2 photopigment removed:

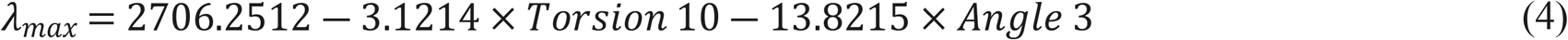
iv. With the guppy Sws2 photopigment removed:

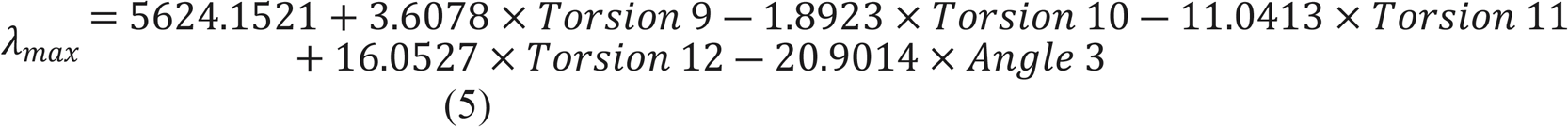

Performance of the “leave-one-species-out” models varied with R^2^ values of 0.84 to 0.93 for the full model and 0.76 to 0.94 for the “leave-one-photopigment-out” approach (S3 Fig). This indicates that our approach generates models with good predictive power for new Sws2 sequences with unknown λ_max_ values that are not included in the model selection process. Furthermore, Angle 3 was present in all models (equations 2–5), as well as in the full model (equation 1), and had the largest weighting within each approach, with the exception of the model where the spotted flounder Sws2 modeling experiment was omitted. In this latter model, Angle 1 was also present and had the largest weighting. In general, these findings imply that Angle 3 (C3–C7–C8) is a significant predictor of λ_max_ values for Sws2 photopigments.

To explain the mechanistic role of 11 putative spectral tuning sites at positions 41, 47, 94, 97, 99, 109, 116, 168, 183, 269, 299 (S1 Fig) revealed by comparative analyses of Sws2a vs. Sws2b sequences [22,23] on the conformation of the chromophore, as predicted by Cortesi et al. (2015) [22], we carried out MD trajectory analyses using visual molecular dynamics (VMD) via visual inspection. It should be noted that except for positions 94 and 269, the other nine putative tuning sites are located far from the chromophore binding pocket. Specifically, position 94 is located in close proximity to the heavy atoms (CE, N+) (Fig 3A) of the K296 residue that is covalently bound to the chromophore, so this site was analyzed in detail. In teleost Sws2b photopigments, position 94 is occupied by a cysteine (C) residue and its sidechain invades the space next to atoms (CE, N+) of K296, resulting in reduced space available for fluctuations of dihedrals/angles of the chromophore. By contrast, position 94 in Sws2a photopigments is occupied by threonine (T), alanine (A) or glycine (G). The T94 residue interacts with S186 and affects the atoms (CE, N+) of the chromophore bound to K296, while A94 and G94 lack large sidechains, thus providing space for the fluctuation of these atoms (CE, N+). To understand the influence of the spectral tuning site at position 94 on the fluctuation of these heavy atoms, we analyzed the dihedrals and geometric bond angles formed by (CE, N+) atoms of K296. We found Angle 2 to be clearly distinguishable for Sws2a compared to Sws2b photopigments (S4 Fig), such that shorter wavelength-shifted (i.e. violet) Sws2b photopigments have wider distributions of Angle 2 compared to longer wavelength-shifted (i.e. blue) Sws2a photopigments.

The spectral tuning site at position 269, as described by Kasagi et al. (2018) [23], is occupied by alanine in all of the Sws2 opsin proteins analyzed in this study, except for *V. variegatus* Sws2a, in which this site is occupied by threonine that results in a shift of the λ_max_ value to longer wavelengths. Analysis of MD trajectories reveals that position 269 is located within the chromophore binding pocket and directly interacts with the β-ionone ring of the chromophore. The methyl group of the A269 side chain forms favorable hydrophobic contacts with the β-ionone ring of the chromophore, but with an A269T substitution, the hydroxyl group of threonine results in the β-ionone ring of the chromophore being positioned more distantly. Among all the 19 structural parameters analyzed that are associated with the chromophore, only Torsions 11 and 12 display spatial relationships such that they could be affected by known tuning sites (see S1 Fig for positions of known tuning sites). Interestingly, Torsion 12 is affected by A269T and was indeed useful for predicting λ_max_ values as can be seen in the combined full model.

## Discussion

We have developed a new model for the prediction of the spectral peaks of absorbance for teleost Sws2 cone photopigments, with high accuracy over a wide range of λ_max_ values (i.e. 397-485 nm). Our approach is based on a predictive model as described in our previous studies [10,34]. In the present study, this approach required Sws2 opsin protein sequence data as input and a known template photopigment structure to build the homology models. MD simulations were then performed on these opsin homology models, with parameters that describe the conformational change in both opsin structure and the covalently-bound chromophore being extracted. From this, a statistical model was built. Similar to our previous studies, our current approach revealed that the structural features of the chromophore and its lysine attachment site play important roles in the prediction of λ_max_ values. In fact, the final predictive model consisted of three terms associated with chromophore conformation. This simple first-order regression model was found to be sufficient to estimate the λ_max_ values of Sws2 photopigments with high accuracy. This study further highlights the versatility of our approach in reliably predicting the λ_max_ values of evolutionarily more distant Sws2 cone opsin sequences, with ~48-51% sequence identity to the bovine RH1 template.

A number of molecular and evolutionary approaches have been used in the field of visual neuroscience to predict visual photopigment λ_max_ values. One common strategy is the application of site-directed mutagenesis to opsin sequences derived directly found to be expressed in a particular extant species, where amino acid substitutions are made followed by measuring the spectral characteristics to identify potential contributions of specific amino acid residues to any observed spectral shifts [2,23,25–27,30]. A similar strategy is to infer the amino acid sequence of the ancestral opsin sequence within a clade, followed by the same technique to experimentally determine λ_max_ values [9,25]. Whereas the latter approach generally involves investigating multiple amino acid substitutions that may or may not be directly related to spectral tuning, the former technique frequently only studies single residue differences. Nonetheless, these methods are frequently used together and have complemented any comparative analyses that preliminarily identify residues that are likely to influence the λ_max_ value of a particular photopigment (e.g. Cortesi et al. (2015) [22]; reviewed by Shichida and Matsuyama (2009) [43]). For example, such an approach identified that a spectral shift of the λ_max_ value of teleost Rh2 photopigments, specifically with sensitivity in the green region of the visible spectrum to blue, was largely due to an E122Q substitution [24,25]. Apart from some vertebrate LWS photopigments [32] and UV-sensitive SWS1 photopigments [26,33], the prediction of other cone opsin λ_max_ values (i.e. SWS2 and RH2 photopigment classes) by manual interrogation of the amino acid sequences alone is extremely difficult and often inaccurate [1,2]. Experimental spectral analyzes by MSP and/or *in vitro* regeneration of photopigments are not always viable options due to the financial limitations and the demand for specialist technical expertise. As such, alternative methods of accurately predicting λ_max_ values to understand photopigment function and ecological adaptation are critical. Our alternative molecular modeling-based approach is, by contrast, simple, accurate and efficient, and does not require site by site substitutions followed by *in vitro* experimentation or complex quantum calculations. Previously, we successfully used a similar approach presented in this study to accurately predicted a broad range of λ_max_ values (467-528 nm) for teleost Rh2 cone photopigments [34] and rod (Rh1) photopigments (444-519 nm) [10]. With this current investigation, our approach now provides accurate predictions of λ_max_ values for Sws2 photopigments from 397-485 nm (i.e. those sensitive to violet vs. blue wavelengths). Not only is our approach useful for accurately predicting λ_max_ values using known spectral tuning sites, it may be used to identify and assess the spectral effects of putative unknown residues on the spectral properties of photopigments. It should be noted, however, that the predictions made using our modeling approach results in λ_max_ values that are based on rhodopsins that utilize a vitamin-A_1_-derived chromophore. This is also the case for experimental approaches using *in vitro* regeneration protocols [2,23,25–27,30]. In some vertebrates (e.g. lampreys [31,44,45], many freshwater teleosts [46], lungfishes [47,48], the green anole lizard *Anolis carolinensis* [49,50]; reviewed in [1–3,51], the visual system is based on porphyropsins that incorporate a vitamin-A_2_-derived chromophore or a combination of both rhodopsins and porphyropsins. Therefore, within the context of biological relevance, the predicted λ_max_ values that result using our approach may have to be converted, where appropriate, to longer wavelengths to account for the possession of a vitamin-A_2_-derived chromophore in native photopigments. This is easily conducted by using a number of rhodopsin-to-porphyropsin transformation algorithms, such as those by Loew and Dartnell [52], Harosi [53], and Whitmore and Bowmaker [14].

One of the key elements of our predictive model is Angle 3 (C3–C7–C8), which is the dominant parameter in predicting the λ_max_ values of Sws2 photopigments. Specifically, our results show that larger values of Angle 3 lead to greater shorter wavelength shifts of λ_max_ values to the violet region of the visible spectrum. Furthermore, Angle 3 is an important parameter when the “leave-one-out” approach was applied at a species level for predicting the spectral peaks of absorbance for unknown opsins, suggesting this parameter is broadly predictive. Other parameters affecting the prediction of λ_max_ values are Torsion 3 and Torsion 12, but the magnitude of their effects is reduced in comparison with that of Angle 3. Nonetheless, like Angle 3, larger values of Torsion 3 and Torsion 12 also short-wavelength-shifted the λ_max_ value. Interestingly, Torsion 12 was the only element of the “three-term” model that is likely to be directly influenced by a known or suspected tuning site (i.e. residue 269). These results explain the possible mechanism that causes the observed long-wavelength shift in the λ_max_ value of the spotted flounder Sws2a photopigment. With the exception of this specific example, other residues affecting Angle 3, Torsion 3, and Torsion 12 cannot be pinpointed to any one particular known or suspected tuning site. Instead, it is likely that multiple tuning sites collectively, either directly or indirectly, influence these chromophore structural features. This finding underscores the power of the homology modeling/MD approach, which takes into consideration the collective influence of the entire amino acid sequence to predict λ_max_ values rather than the sole specific contributions of individual sites.

Within the chromophore of the *D. rerio* (zebrafish) Sws2 photopigment, Torsion 3 has a relatively higher median value compared to the median value range of Torsion 3 of Sws2 photopigments found in other species, while Torsion 12 has a relatively lower median value compared to the median value range of Torsion 12 of other Sws2 photopigments. These “out-of-range” median values of Torsion 3 and Torsion 12 lead to a less accurate prediction of the λ_max_ value for the *D. rerio* Sws2 photopigment. Based on the evolutionary relationships of teleost Sws2 opsin protein sequences examined in this study, it appears that *D. rerio* (and maybe other cyprinids) diverged from the main *sws2* opsin clade before *sws2* duplicated into *sws2a* and *sws2b* subclasses. It is possible, therefore, that the zebrafish Sws2 opsin protein holds the retinal chromophore in a distinct conformation vs. the other Sws2 proteins investigated in this study, but which also results in a λ_max_ value that is similar to that exhibited by the Sws2b photopigment subclass. Thus, teleost Sws2 photopigments may have independently evolved more than one opsin-chromophore conformation strategy for attaining short-wavelength shifts of the λ_max_ value to the violet region of the visible spectrum. Such knowledge means that the model presented here and its predictive power might be improved in the future if a more diverse set of Sws2 photopigments is used to develop a more all-encompassing Sws2 model and/or to develop distinctive models for some selected phylogenetic groups.

Comparative analyzes of Sws2a vs. Sws2b photopigments have revealed candidate spectral tuning sites that could potentially explain the sensitivity to violet vs. blue wavelengths for Sws2b and Sws2s subgroup, respectively [22]. Thus, MD simulations can serve as a valuable, complementary tool to understand the contributions of candidate tuning sites to spectral shifts in the λ_max_ value. From MD trajectory analysis, Angle 2 was identified as a potential parameter that is affected by the presence of different residues at spectral tuning site 94. However, we note that as Angle 2 did not display a significant linear correlation with the actual λ_max_ value, Angle 2 was not considered as a candidate for the model selection procedure; as such, Angle 2 did not appear in the predictive “three-term” model.

In conclusion, we have successfully tested our previously studied molecular modeling approach combining homology modeling, MD simulations and structural information of chromophore conformations and to accurately predicted the λ_max_ of Sws2 opsins. In future studies, we plan to consider additional features of each parameter (e.g. narrow vs. broad distribution) other than simple linear correlations, which may further improve the predictive power of the resulting models. We will also expand our approach to extensively study the more phylogenetically distant classes of opsins, with large number of opsins included in the dataset. Finally, we aim to develop model(s) using a dataset of individual distinct classes of opsins to generate an online web platform for accurately predicting the λ_max_ values for any unknown vertebrate photopigment.

## Methods

### Phylogenetic analysis

The alignment and phylogenetic reconstructions were performed using the function “build” of ETE3 v3.1.1 [54] as implemented on the GenomeNet (https://www.genome.jp/tools/ete/). The multiple sequence alignment was provided as input file. ML tree was inferred using PhyML v20160115 ran with model and parameters: --alpha e --pinv e -f m -o tlr --bootstrap -2 --nclasses 4 [55]. Branch supports are the Chi2-based parametric values return by the approximate likelihood ratio test.

### Homology modeling

Eleven teleost Sws2 cone opsin amino acid sequences (Table 1), namely *V. variegatus* (spotted flounder) [23], *M. achne* (slime flounder) [23], *P. olivaceus* (olive flounder) [23], *P. reticulata* (guppy) [39], *O. latipes* (Japanese rice fish; medaka) [20], *L. goodei* (bluefin killifish) [40], and *D. rerio* (zebrafish) [1], were downloaded from the UniProt database (https://www.uniprot.org/). These were selected because the corresponding spectral peaks of absorbance for their Sws2 photopigments (when reconstituted with a 11-*cis* retinal chromophore) have been experimentally measured. Collectively, these photopigments exhibit a wide range of λ_max_ values from 397-485 nm. An experimental 3D structure of a Sws2 cone photopigment is not available; hence, to build a homology model of 11 Sws2 photopigments, a template search was carried out using SWISS-MODEL (https://swissmodel.expasy.org/). The closest homologue (~50% sequence identity) with a high-quality 3D structure was found to be that of the bovine rod opsin (RH1). A high-resolution crystallographic structure of the bovine RH1 photopigment (PDB ID 1U19, 2.2 Å) [38], which lacks mutations and has an 11-*cis* retinal chromophore covalently bound within its binding pocket, was downloaded from the Protein Data Bank. 3D coordinates of the bovine rod opsin structure were then used to build the homology models of 11 teleost Sws2 cone opsin sequences. The structure prediction wizard from the PRIME module of the Schrödinger suite was used for building a homology model for each protein sequence [56,57]. The non-templated loops were refined using the refine loops module of PRIME and a generated model structure was validated by generating a Ramachandran plot by analyzing acceptable phi-psi regions of residues. The final homology model was modified to remove the intracellular unstructured coil (~25 residues) region towards the carboxyl-terminus to prevent it from crossing the periodic boundaries during the molecular dynamics (MD) simulation.

### Molecular dynamics (MD) simulation

All 11 Sws2 photopigment homology models were subjected to atomistic MD simulations using our protocol for input file generation and the system setup for MD simulations reported in our previous studies [10,34]. Briefly, the homology models of each Sws2 opsin sequence with the chromophore bound covalently to the lysine residue in the binding pocket (K296) were uploaded to the CHARMM-GUI webserver (http://charmm-gui.org). Each system was placed in lipid bilayers and hydrated using a hexagonal solvent box with a 15 Å TIP3P water layer. The charge of the system was neutralized with 150 mM NaCl. The CHARMM36m forcefield [32] parameters were selected for all the components of the systems. After minimization and short equilibration simulations with harmonic restraints, the systems were subjected to 100 ns atomistic MD simulations. The production simulations were performed under an NPT ensemble for 100 ns using a Parrinello-Rahman barostat [58] with semi-isotropic pressure coupling and a Nosѐ-Hoover thermostat [59]. During the production MD simulations, snapshots were saved every 10 ps. GROMACS-2018.3 [60] was used for all 11 MD simulations. The visualization and analysis of MD trajectories were carried out using Visual Molecular Dynamics package [61]. GROMACS trajectory analysis tools were used to analyze Root Mean Square Fluctuations (RMSF) and 19 different internal degrees of freedom of the chromophore (i.e. torsion angles and geometric bond angles).

### Quantification and statistical analysis

To determine the best linear regression model for predicting Sws2 λ_max_ values, we curated a shortlist of structural parameters associated with the chromophore based on their linear correlation to the experimental λ_max_ value for each photopigment. (S2 Fig) From our list of 19 parameters, a shortlist was generated that composed of the medians of Torsion 3, Torsion 9, Torsion 10, Torsion 11, Torsion 12, and geometric Angles 1 and 3. A model selection procedure was then performed using the regsubsets function of the “leaps” R library (https://cran.r-project.org/web/packages/leaps/leaps.pdf). This evaluated all possible model subsets using the shortlisted angles and ranked them according to their Bayesian Information Criterion (BIC) value [62], which quantifies the explanatory power of a model with a penalty for the number of terms included. The resulting best-fit model was further evaluated via a “leave-one-out” procedure by reweighting the parameters of the best-fit model after removing each photopigment iteratively. For each Sws2 photopigment stimulation, the parameters of the best-fit model were reweighted with the given Sws2 photopigment data removed. This reweighted model was subsequently used to predict the spectral peak of absorbance for the omitted Sws2 opsin sequence.

Additionally, to further validate our approach and to simulate performance at predicting the spectral peaks of absorbance for Sws2 photopigments with unknown λ_max_ values, a systemic “leave-one-species-out” process was also performed. For a given species, both Sws2a and Sws2b opsin sequences were removed from the dataset. A new model was chosen using the full model selection procedure detailed above. Since removing two opsins results in a smaller sample of data, overfitting (i.e. a model with the same number of terms as the number of datapoints it is being fit to) is a possible risk factor. To mitigate this, the generated model space was explored within +2 BIC value of the best BIC ranked value. The resulting model was then used to predict the spectral peak of absorbance for the removed Sws2a and Sws2b photopigments.

## Acknowledgements

We thank Dr. F. Marty Ytreberg for constructive feedback on an earlier version of the manuscript.

## Supporting information captions

**S1 Fig. Sequence alignment of Sws2 opsins used in the simulations, together with the bovine rod opsin template**. Red arrows indicate amino acid positions (using numbering standardized to bovine rod opsin) highlighted by Cortesi et al. (2015) [22] that were identified as likely to have contributed to the functional diversification of teleost Sws2 opsin sequences, particularly those that distinguish Sws2b photopigments (violet-sensitive) from the more long-wavelength-shifted Sws2a photopigments (blue-sensitive). The blue arrow indicates position 269, that is likely to be the spectral tuning site that mediates a shift in the λ_max_ value of *V. variegatus* Sws2a to 485 nm [23]. A large orange asterisk (*) depicts a conserved lysine (K) residue at position 296 that is required for the formation of the Schiff-base linkage to the retinal chromophore. The gray bars below each alignment indicate a quality score, such that lower scores correspond to greater amino acid variability in the column. All species and photopigment subclasses are followed by experimentally measured λ_max_ values.

**S2 Fig. Frequency distribution of all analyzed torsions and geometric angles observed for each photopigment simulation.** Each panel represents the frequency distribution of an individual torsions and angles associated with photopigment. Blue and violet colors indicate the relative spectral peaks of absorbance of each Sws2 photopigment, corresponding to λ_max_ values >430 nm vs. λ_max_ values <430 nm, respectively.

**S3 Fig. Experimental spectral peaks of absorbance (i.e. the λ_max_ value) compared to predicted λ_max_ values generated by the model resulting from “leave-one-system-out” analysis.** The prediction for the removed Sws2a and Sws2b photopigments are indicated by blue and violet squares, respectively. Black squares correspond to the Sws2 photopigments used in the model selection procedure with the black line indicating the R^2^ for the model. Red circles and dashed line correspond to the leave-one-out analysis for single photopigments included in the model. The grey line corresponds to a 1:1 relationship.

**S4 Fig. Frequency distribution of Angle 2 observed in Sws2a and Sws2b of each species used in the simulation.** Each panel represents the frequency distribution of Angle 2 observed in individual species with Sws2a and Sws2b (*L. goodei, O. latipes, P. reticulata, V. variegatus*). Sws2a and Sws2b photopigments are indicated by blue and violet colors, respectively.

**S1 Movie. Molecular dynamics simulations of Sws2a opsin with 11-*cis retinal from V. variegatus* (spotted flounder) embedded in the lipid bilayer and water.**

